# Macrophage PHOX Restricts *Aspergillus fumigatus* Germination and Hyphal Development Through ROS-Dependent Immunity

**DOI:** 10.64898/2026.10.13.758795

**Authors:** M. Lynette Goins, Sourabh Dhingra, Emily E Rosowski

## Abstract

*Aspergillus fumigatus* is a fungal pathogen that causes invasive disease in immunocompromised individuals. Innate immune cells, including macrophages and neutrophils, are essential for controlling *A. fumigatus*, but the mechanisms by which specific phagocytes control distinct stages of infection remain incompletely understood. In particular, the contribution of macrophage-derived reactive oxygen species (ROS) to conidial clearance and control of fungal germination and hyphae remains unclear. Here, we used larval zebrafish with mutation of *p22*, an essential subunit of the phagocyte NADPH oxidase (PHOX) complex, and neutrophil-deficiency, to define the role of macrophage-derived ROS during *A. fumigatus* infection. Mutation of *p22* or pharmacological ROS scavenging increased host susceptibility, supporting a role for ROS in macrophage-mediated defense. However, macrophage ROS restricts fungal developmental progression rather than promoting clearance of conidia. Using Δ*pksP* conidia, we confirmed that fungal melanin limits the localization of p22 to phagosomes, but found no difference in conidial killing of Δ*pksP* conidia, supporting the finding that PHOX-generated ROS are not required for macrophage conidial killing. However, Δ*pksP* conidia have defective virulence only in ROS-producing larvae, as melanin inhibits ROS-mediated control of germination and hyphal growth. These findings identify macrophage PHOX activity as a stage-specific regulator of *A. fumigatus* infection that primarily restricts fungal developmental progression, while PksP-produced melanin promotes immune evasion by limiting macrophage ROS-dependent antifungal activity. These findings provide insight into how macrophages inhibit the transition from conidia to invasive fungal growth and how *A. fumigatus* evades this defense, with implications for understanding determinants of disease progression during invasive aspergillosis.

## Introduction

*Aspergillus fumigatus* (*A. fumigatus*) is a ubiquitous filamentous fungus and major airborne pathogen responsible for invasive aspergillosis, a disease affecting over 250,000 to 300,000 individuals annually (Bongomin et al., 2017; Fisher et al., 2020). In immunocompromised individuals, inhaled conidia can germinate into invasive hyphae within the respiratory tract, leading to severe and often disseminated disease (Cadena et al., 2016; Latgé & Chamilos, 2019). In healthy individuals, innate immune cells provide the first line of defense against *A. fumigatus*. Macrophages and neutrophils recognize fungal cell-wall components that become exposed during conidial swelling and germination (Aimanianda et al., 2009; Hohl et al., 2005). Traditionally, macrophages are thought to primarily control conidia, while neutrophils restrict hyphal growth (Hohl & Feldmesser, 2007; Latgé & Chamilos, 2019; Schaffner et al., 1982).

However, macrophages can exhibit broad antifungal functions, including phagocytosis and killing of conidia, inhibition of germination, and restriction of hyphal growth (Cornish et al., 2008; Latgé & Chamilos, 2019; Philippe et al., 2003; Rosowski et al., 2018). Macrophages have several mechanisms for controlling *A. fumigatus*, including phagosomal acidification and ROS production. Previous work has demonstrated that macrophages generate NADPH phagocyte oxidase (PHOX)-dependent ROS in response to *A. fumigatus* and implicated this pathway in antifungal activity (Philippe et al., 2003). NOX2-derived ROS can also regulate LC3-associated phagocytosis, linking PHOX activity to phagosomal antimicrobial functions (Chamilos et al., 2016; Ligeon et al., 2021).

Phagocyte ROS production through the PHOX complex is an important innate immune function that helps protect against *A. fumigatus* infection. Patients with chronic granulomatous disease (CGD), who lack functional PHOX activity, are highly susceptible to *Aspergillus* infection (Latgé & Chamilos, 2019; Marciano et al., 2015; Winkelstein et al., 2000). However, because PHOX is active in both macrophages and neutrophils, its cell type-specific contributions to antifungal immunity remain unclear. Neutrophil PHOX activity has been implicated in limiting fungal expansion after germination and regulating inflammatory responses, but it appears to have little effect on initial germination (Grimm et al., 2013; Schoen et al., 2020). Evidence regarding macrophage PHOX activity is less consistent. Alveolar macrophages can inhibit conidial germination in the absence of detectable PHOX activity, suggesting that this early response may be PHOX-independent (Cornish et al., 2008). In contrast, ROS production by alveolar macrophages has been reported to be necessary for killing conidia after phagocytosis (Philippe et al., 2003). Furthermore, genetic studies in mice have implicated macrophage PHOX activity in limiting fungal germination and protecting against pulmonary aspergillosis (Grimm et al., 2013). Together, these findings suggest that PHOX contributes to macrophage-mediated antifungal defense, but that its precise function may vary according to the stage of fungal development and the antifungal outcome examined.

Also complicating this question is the ability of *A. fumigatus* to evade macrophage antimicrobial defenses through melanin produced via the *pksP* pathway. PksP-produced melanin reduces phagolysosome maturation and intracellular killing of conidia by macrophages (Jahn et al., 2002) and can inhibit NADPH oxidase-dependent responses by excluding the p22 subunit from the phagosome (Akoumianaki et al., 2016; Chamilos et al., 2016). How this immune evasion strategy affects macrophage control across distinct stages of fungal development and influences morbidity and mortality of infected hosts remains unclear. In mammalian models, it is difficult to disentangle whether PHOX is required for spore killing, inhibition of germination, or restriction of the transition to invasive hyphae, as these processes cannot be directly observed over time in vivo.

To overcome this limitation, we use the larval zebrafish host model, which allows macrophage functions including spore killing, inhibition of germination, and restriction of hyphal transition to be directly visualized and quantified over time in living vertebrate animals (Knox et al., 2017; Rosowski et al., 2018). Zebrafish possess a conserved innate immune system with humans and lack an adaptive immune response during early development, enabling specific analysis of innate antifungal mechanisms (Lam et al., 2004; Yoder et al., 2002). The optical transparency of larvae and availability of transgenic fluorescent reporters allow real-time visualization of host-pathogen interactions at single-cell resolution (Knox et al., 2017; Marques et al., 2019; Rosowski, 2020). The larval zebrafish model has been used extensively to define innate immune responses to *Aspergillus* infection, revealing distinct contributions of macrophages and neutrophils to controlling conidial and hyphal stages of fungal infection (Knox et al., 2014), identifying intracellular germination of phagocytosed conidia as a key feature of *A. fumigatus* pathogenesis (Koch et al., 2019), and uncovering host signaling pathways, including cyclooxygenase-dependent PGE2 production and Rac2-mediated macrophage functions, that restrict fungal germination and invasive hyphal development (Tanner & Rosowski, 2024; S. Thrikawala et al., 2022). Together, these studies highlight the utility of the larval zebrafish system for dissecting stage-specific host mechanisms that regulate *Aspergillus* infection.

Here, we use the larval zebrafish model to define the stage-specific role of macrophage PHOX activity during *A. fumigatus* infection through genetic, pharmacological, and live-imaging approaches. We find that macrophage-derived ROS promotes host survival primarily by restricting fungal germination and invasive hyphal growth, rather than by promoting spore killing. We further demonstrate that PksP-produced melanin antagonizes PHOX-dependent macrophage defense, with loss of melanin increasing the contribution of PHOX to conidial clearance and altering the control of fungal developmental progression. Together, these findings establish macrophage PHOX activity as an important mechanism for controlling fungal developmental progression and suggest that PksP-mediated modulation of ROS-dependent immunity contributes to *A. fumigatus* virulence.

## Methods

### Zebrafish lines and maintenance

All experimental procedures of zebrafish embryos and larvae were performed, and adult zebrafish were maintained and handled, according to protocols approved by the Clemson University Institutional Animal Care and Use Committee (AUP2024-0270, AUP2025-0094, AUP2025-0095). Adult and larval zebrafish were maintained as previously described (Thrikawala et al., 2022). Embryos/larvae were anesthetized using buffered tricaine prior to any experimental procedures. Larvae were euthanized at 4°C, and adults were euthanized with buffered tricaine. Mutant and transgenic strains used are listed in Table 1 and were maintained in the AB genetic background. For experiments with a *p22 (cyba)* mutant line (sa11798), which has previously been validated to not express p22 protein (Schoen et al., 2020), heterozygote adults were in-crossed and larvae were genotyped at the conclusion of the experiment by isolating genomic DNA from whole larvae in 10 mM NaOH at 95°C for 20 minutes and dCAPS PCR and RFLP digestion with MseI (NEB) as previously described (Schoen et al., 2020), (F: *sa11798_dCAPS_F* 5’ CTTTTGGACCCCTGACCAGAAATTA 3’; R: *sa11798_dCAPS_R* 5’ TGGCTAACATGAACCCTCCA 3’). For experiments with transgenic fluorescent lines, embryos were screened for fluorescence on a SteREO Discovery.V12 fluorescent microscope (Zeiss), at 2 days post fertilization (dpf), prior to experimentation.

**Table 1.** Zebrafish lines used in this study.

| Line | Purpose | Reference |
| --- | --- | --- |
| *mfap4:BFP* | express BFP in macrophage cytosol | (Rosowski et al., 2018) |
| *mpx:mCherry-2A-rac2D57N* | neutrophil-defective | (Deng et al., 2011) |
| *mfap4:p22-TagRFPT__cryaa:mCherry* | express fluorescently-tagged p22 in macrophages | This study |
| *p22 sa11798* | mutant line with premature stop codon | (Schoen et al., 2020) |

### Generation of transgenic zebrafish with fluorescently-tagged p22

Coding sequence for *p22*, without a stop codon, was amplified with Q5 polymerase (NEB) from cDNA made with iScript RT supermix (Bio-Rad) from total RNA isolated from 2-3 dpf zebrafish larvae with TRIzol (Invitrogen) using F primer: 5’-CTCAAGTGATCATGGCGAAG-3’ and R primer: 5’-CTGTTACTTGTTGCCATCACCT-3’. Purified PCR product was then used as template for a second PCR using F primer: 5’-ATGACTTTAGATCGTGGTCGAGGATCCATGGCGAAGATTGAGTGG-3’ and R primer: 5’-CACCATTCCGCTTCCGCTTCCGGATCCTTCGTTGATGGTGACAGA-3’ to add overlaps for HiFi cloning (NEB) into BamHI site of a previously-generated N-terminal tagging vector (Thrikawala et al., 2026), to generate Tol2-*mfap4:p22-TagRFPT cryaa:mCherry*. This plasmid was co-injected with Tol2 *transposase* mRNA into 1 cell embryos as previously described (Thrikawala et al., 2023). Resulting embryos/larvae were screened for mCherry expression in the lens of the eye, grown to adulthood, and out-crossed to identify a germline founder.

#### CRISPR-Cas9 targeting of p22

Guide RNAs were designed and injected as previously described (Thrikawala et al., 2023). For p22 targeting, two guide RNAs were designed to induce double-stranded breaks within exon 4 and exon 5 of the target gene (Supp Fig 1). The gRNA sequences were as follows: p22_gRNA_1: 5′-ACGTAATAGTTTCTGGTCAG-3′; p22_gRNA_2: 5′-TGGCTAACATGAACCCTCCA-3′. Guide RNAs were *in* vitro transcribed and co-injected with Cas9 protein (PNA Bio NC1279639) into one-cell stage zebrafish embryos, as previously described (Thrikawala et al., 2022) to generate *p22* crispants which were used directly for experiments.

To assess CRISPR/Cas9-mediated editing, PCR primers were designed to amplify the genomic region spanning the two p22 gRNA target sites (Supplemental Figure 1). The forward primer, p22_gf1 (5’-CAGCGGTCAGTACTGCTTTACA-3’), was located upstream of the first gRNA target site, and the reverse primer, p22_gr2 (5’-GGTAGATCAAACTTGCCATTCC-3’), was located downstream of the second gRNA target site. This primer pair amplified the region encompassing both target sites, allowing detection of excision of the intervening DNA. The expected sizes of the intact and excised amplicons were approximately 300 bp and 100 bp, respectively.

As a control, guide RNAs targeting luciferase were used in parallel with the experimental gRNAs, as previously described (Thrikawala et al., 2023). The luciferase gRNA sequences were as follows: luciferase_gRNA1: 5′-TTGGAAACGAACACCACGGT-3′; luciferase_gRNA2: 5′-ACAACTTTACCGACCGCGCC-3′.

### *Aspergillus fumigatus* Spore Preparation and Infection of Zebrafish Larvae

Preparation and injection of spores into larvae was performed as previously described (Knox et al., 2014; Thrikawala et al., 2022; Thrikawala & Rosowski, 2020)*. A. fumigatus* spores were plated at 1 × 10⁶ spores on glucose minimal media (GMM) plates and incubated for 3 days at 37°C. Spores were collected, washed with 0.01% Tween, filtered through Miracloth (MilliporeSigma; EM475855-1R), pelleted by centrifugation at 900 × g for 10 min, washed in sterile phosphate-buffered saline (PBS), pelleted again, and resuspended in 5 mL sterile PBS. The suspension was filtered again through Miracloth, and spore concentration was determined using a hemocytometer. Spores were diluted to 1.5 × 10⁸ spores/mL in sterile PBS for injection and kept at 4°C for up to 6 weeks. Before injection, 2 dpf larvae were anesthetized in E3 containing 0.3 mg/mL buffered tricaine.

Larvae were microinjected in the hindbrain ventricle with a spore suspension prepared in PBS containing phenol red using a microinjection system (Applied Scientific Instrumentation) equipped with a pressure injector, micromanipulator, micropipette holder, footswitch, and back-pressure unit. The injection volume was adjusted to deliver approximately 40–50 spores per larva, consistent with previously described infection protocols (Thrikawala & Rosowski, 2020). Following injection, larvae were washed with fresh E3 medium. *Aspergillus fumigatus* strains used in this study are listed in Table 2.

**Table 2.**
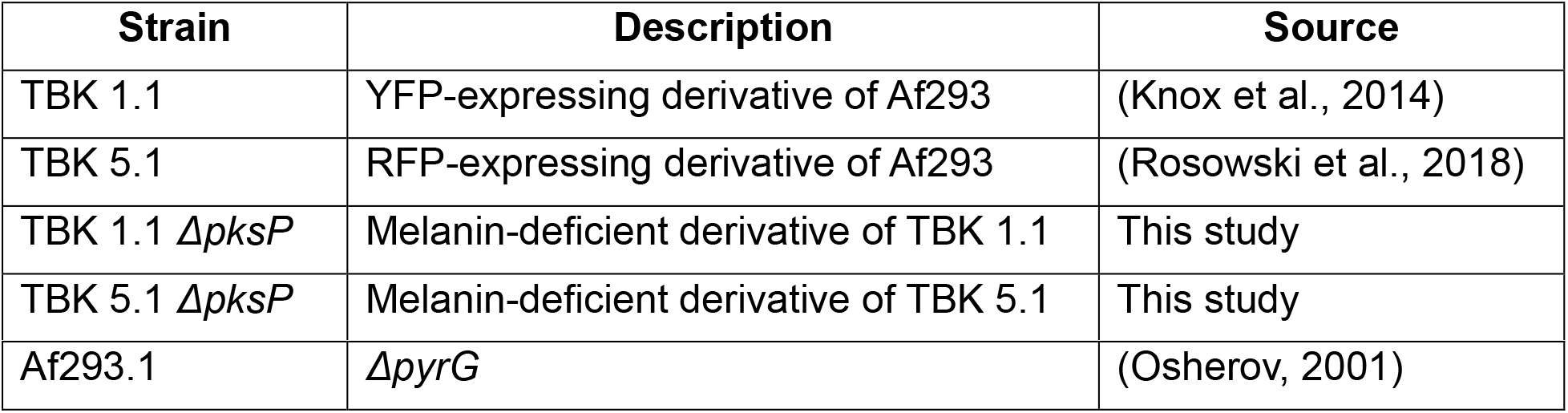
*Aspergillus fumigatus* strains in this study.

### Generation of TBK 1.1 *ΔpksP* and TBK 5.1 *ΔpksP*

The conidial pigment polyketide synthase gene, *pksP* (*Afu2g17600*), was deleted from the YFP-expressing strain TBK1.1 (Knox et al., 2014) and the mCherry-expressing strain TBK5.1 (Rosowski et al., 2018) using CRISPR-Cas9. Briefly, the homology-directed repair cassette was amplified from plasmid pBC-Hygro (FGSC) using primers SD720 and SD721. The crRNAs targeting the *pksP* locus were ordered from IDT. Protoplast-mediated transformation was performed as previously described (Al Abdallah et al., 2017). White colonies were selected and deletion of *pksP* was confirmed by Southern blot. Primer and crRNA sequences were as follows: SD720: 5′-TCTTTTCAAAGTGGGCTGTTGTTTTCTTCTGCTTTACCGGTCGCCTCAAACAATGCTCT-3′; SD721: 5′-CACAACTCCACCCCCAAACCTAGGAACTCACCCCAGTCTGAGAGGAGGCACTGAT GCG-3′; pksP guide F: /AlTR1/rArUrArCrCrArArArArArArCrCrCrCrArUrGrUrGrGrUrUrUrUrArGrArGrCrUrArUrG rCrU/AlTR2/; and pksP guide R: /AlTR1/rCrCrArCrGrGrCrCrArUrGrArGrUrUrCrCrUrArGrGrUrUrUrUrArGrArGrCrUrArUr GrCrU/AlTR2/.

### CFU Enumeration

CFU enumeration was performed as previously described (Rosowski et al., 2018). Individual larvae were homogenized and plated on GMM agar, and plates were incubated at 37°C for 3 days before colony enumeration. Larvae infected with the Af293.1 (Δ*pyrG*) *A. fumigatus* strain were plated on GMM agar supplemented with uracil and uridine.

### Survival Assays

Eight larvae from each replicate were homogenized immediately after infection and plated on GMM agar to quantify the inoculum by CFUs. The average measured inoculum for each experiment is reported in the corresponding figure legend. The remaining infected larvae were transferred to fresh E3 medium, individually placed into 96-well plates, and monitored daily for survival over a 7-day period. Larvae were scored as dead based on the absence of a heartbeat.

### Live Imaging of Infection

For imaging experiments, zebrafish larvae were continuously treated with 100 µM N-phenylthiourea (PTU) starting at 1 dpf to inhibit melanization. Following infection, larvae were transferred to 48-well plates.

Daily imaging was performed as previously described (Thrikawala & Rosowski, 2020). On each day of live imaging, larvae were anesthetized with 0.3 mg/mL buffered tricaine in E3 and placed into a zWEDGI device (Huemer, Squirrell, Swader, LeBert, et al., 2017; Huemer, Squirrell, Swader, Pelkey, et al., 2017) to orient them laterally for imaging of the hindbrain ventricle. Larvae were imaged using a Zeiss Cell Observer Spinning Disk confocal microscope on an Axio Observer 7 microscope stand with a confocal scanhead (Yokogawa CSU-X) and a Photometrics Evolve 512 EMCCD camera. Either a Plan-Apochromat 20x objective (0.8 NA) or an EC Plan-Neofluar 40x objective (0.75 NA) and ZEN software were used to acquire Z-stack images of the hindbrain area. After imaging, larvae were rinsed in E3 and returned to their corresponding well in the 48-well plate with 100 µM PTU.

### NAC Treatment

N-acetyl-L-cysteine (NAC; Sigma-Aldrich, A9165) was dissolved in PBS to prepare a 1 M stock solution and stored at −20°C in aliquots. A fresh 100 mM NAC working solution was prepared from the 1 M stock solution each day and added to fresh E3 medium immediately following infection, and refreshed each day, to achieve a final concentration of 200 µM. Control larvae were maintained in E3 containing an equivalent volume of PBS, alsoreplaced daily throughout the experiment.

### H2DCFDA ROS Staining

For reactive oxygen species (ROS) staining, 2′,7′-dichlorodihydrofluorescein diacetate (H2DCFDA; Invitrogen, D399) was dissolved in DMSO to prepare a 100 µM stock solution and stored at −20°C in aliquots. At 2 dpi, the H2DCFDA stock solution was diluted in filter-sterilized 1% phenol red to a final concentration of 10 µM and microinjected (1–2 nL) into the zebrafish larval hindbrain. Immediately following injection, larvae were fixed in a PIPES-based fixation solution prepared as previously described (Miskolci et al., 2019) using piperazine-N,N′-bis(2-ethanesulfonic acid) (PIPES; Thermo Scientific Chemicals, AC172610250). Fixed larvae were kept in the dark at 4°C and imaged the following day for ROS fluorescence as described in the live-imaging section above.

### Image Analysis

All image analysis was performed using Fiji.

ROS imaging analysis (Figure 1) was performed using a maximum-intensity projection of 10 Z-slices from the central region of the zebrafish hindbrain containing both macrophages and *A. fumigatus*. A region of interest (ROI) was manually drawn on the composite image to encompass the macrophage, fungal, and ROS-associated fluorescence signals. The same ROI was then applied to the ROS channel, and the mean fluorescence intensity was quantified within the defined region.

**Figure 1.**
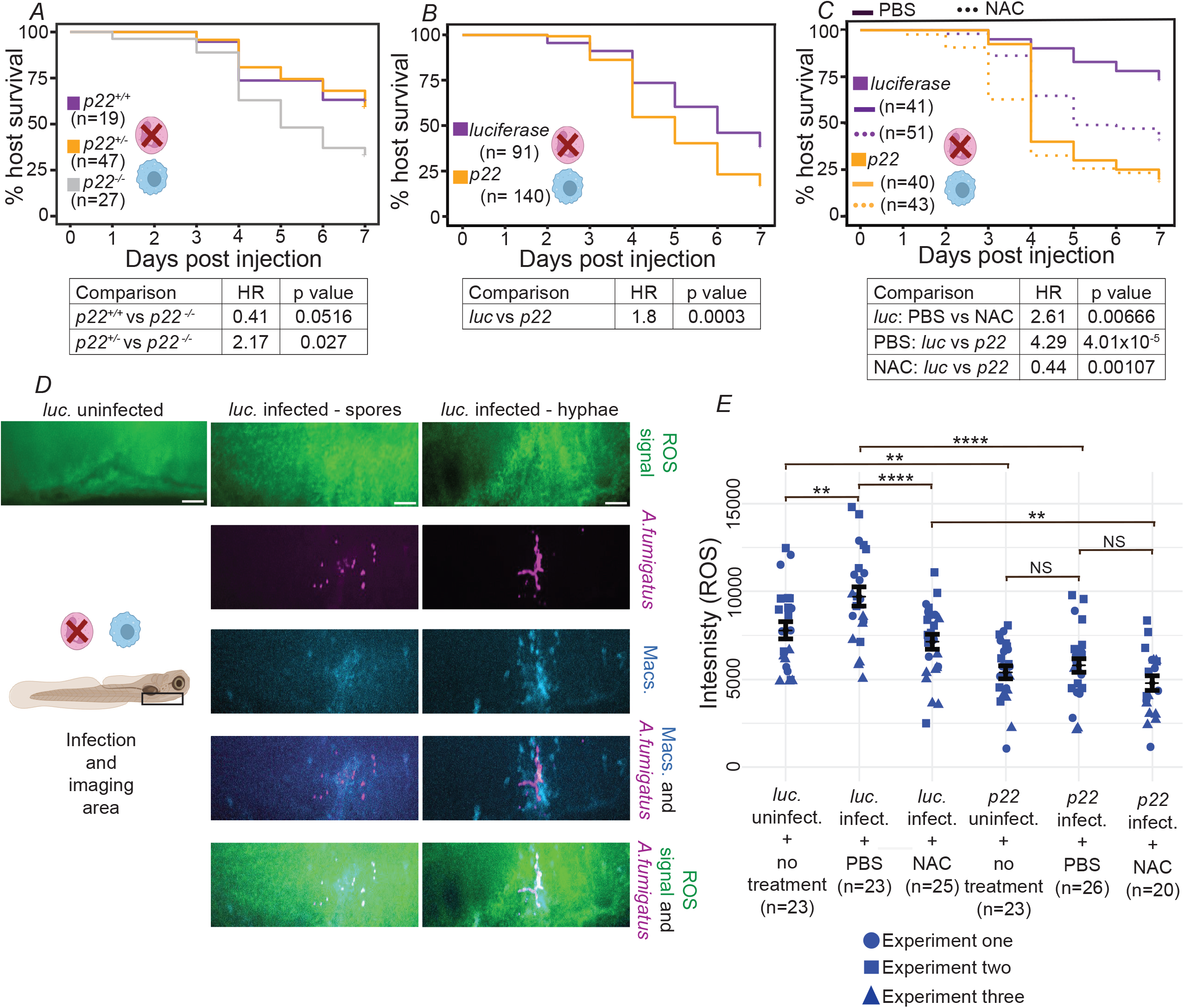
ROS promotes macrophage-mediated protection against *A. fumigatus* infection in larval zebrafish. **A–C.** Survival of neutrophil-defective (*mpx:rac2D57N*) zebrafish larvae following hindbrain ventricle infection at 2 days post-fertilization (dpf) with the YFP-expressing *A. fumigatus* strain TBK1.1. Larvae also carried a stable mutation in *p22* (A) or were *p22* crispants or *luciferase* controls (B, C). Larvae in (C) were also treated with N-acetyl-L-cysteine (NAC; 200 µM) or phosphate-buffered saline (PBS) following infection. Larvae were injected with an average dose of 42 CFUs (A), 51 CFUs (*luc*) or 44 CFUs (*p22*) (B), or 42 CFUs (*luc*) or 39 CFUs (*p22*) (C). Data represent three pooled independent replicates. **D–E.** Neutrophil-defective (*mpx:rac2D57N*) larvae were infected with the RFP-expressing *A. fumigatus* strain TBK5.1 and treated with either N-acetyl-L-cysteine (NAC; 200 µM) or PBS. At 2 days post-infection (dpi), larvae were injected into the hindbrain ventricle with 2′,7′-dichlorodihydrofluorescein diacetate (H2DCFDA) and immediately fixed using a PIPES-based fixation solution. PBS-injected larvae served as a negative control for H2DCFDA staining. (D) Representative images of H2DCFDA fluorescence marking reactive oxygen species (ROS). Scale bar, 20 µm. (E) Quantification of fluorescence intensity. Individual points represent single larvae, with point shape indicating replicate. Data represent three replicates with a total N of 140 larvae and are presented as estimated marginal means (emmeans) ± SEM.

Co-localization analysis (Figures 2 and 4) was performed using Z-stacks of individual larvae. Total fungal spores within each larva were quantified, and spores located within macrophages were identified based on the macrophage and fungal fluorescence signals. Co-localization events were defined as spatial overlap between p22 and spore fluorescence signals within macrophages. For each larva, the total number of spores, the number of intracellular spores, and the number of intracellular spores co-localizing with p22 were quantified. The percentage of intracellular spores co-localizing with p22 was calculated for each larva, and larvae were additionally categorized based on whether at least one intracellular spore co-localized with p22.

**Figure 2.**
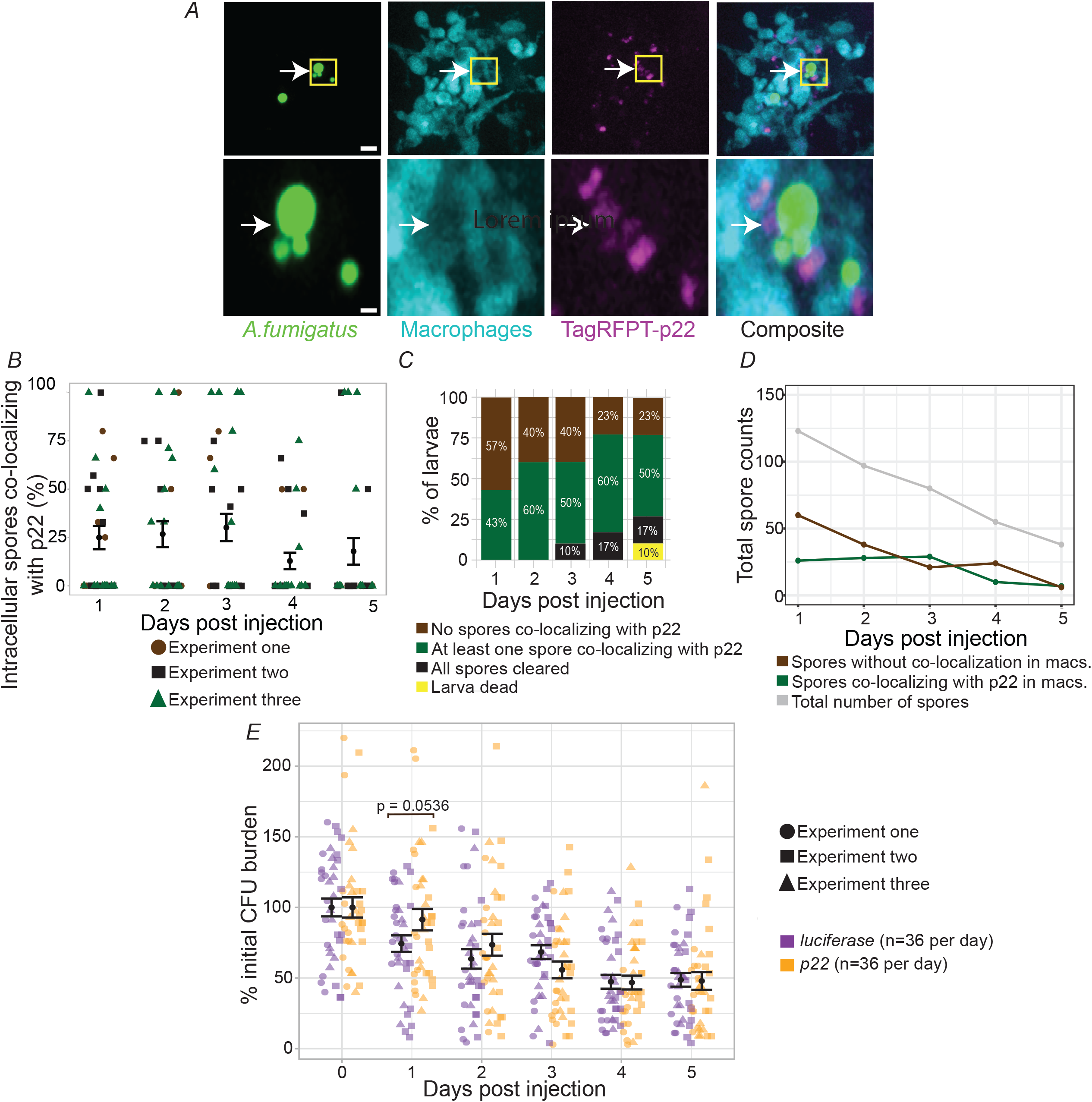
p22 does not play a major role in killing of *A. fumigatus* spores by macrophages. **A-D.***mfap4:BFP* (macrophage-labeled) zebrafish were crossed with *mfap4:TagRFPT-p22* (p22-labeled) zebrafish and larvae were infected with an average of 18 spores of the YFP-expressing *A. fumigatus* strain TBK1.1 and imaged daily from 1-5 dpi. Data shown are pooled from three independent experiments, with a total N of 30 larvae. (A) Representative confocal images showing co-localization of p22 with *A. fumigatus* spores within macrophages. White arrows indicate co-localization events, and yellow boxes indicate zoomed in area. Scale bar, 15 µm. (B) Quantification of the percentage of spores within macrophages that co-localize with p22 at each dpi was quantified. Each dot represents an individual larva. Bars indicate estimated marginal means (emmeans) ± SEM. (C) Quantification of the percentage of larvae containing at least one spore co-localizing with p22 within macrophages at each dpi. (D) Total spore counts in different categories across all larvae**. E.** *p22* crispant or *luciferase* control larvae were generated, infected with a Δ*pyrG* strain of *A. fumigatus*, and homogenized for CFU counts from 0–5 dpi, with a total *N* of 36 larvae per condition per day. CFU counts were normalized to the average injection dose at 0 dpi. Each point represents an individual larva, with point shape indicating replicate. Data represent three independent experiments, with 36 total larvae per dpi per condition, with bars indicate estimated marginal means (emmeans) ± SEM.

Fungal burden and macrophage area (Figure 3) were quantified in individual larvae from maximum-intensity Z-projections generated from Z-stacks encompassing the region of infection. An ROI was manually defined around the infection site, and thresholding was applied to quantify the area of fungal and macrophage fluorescence signals.

**Figure 3.**
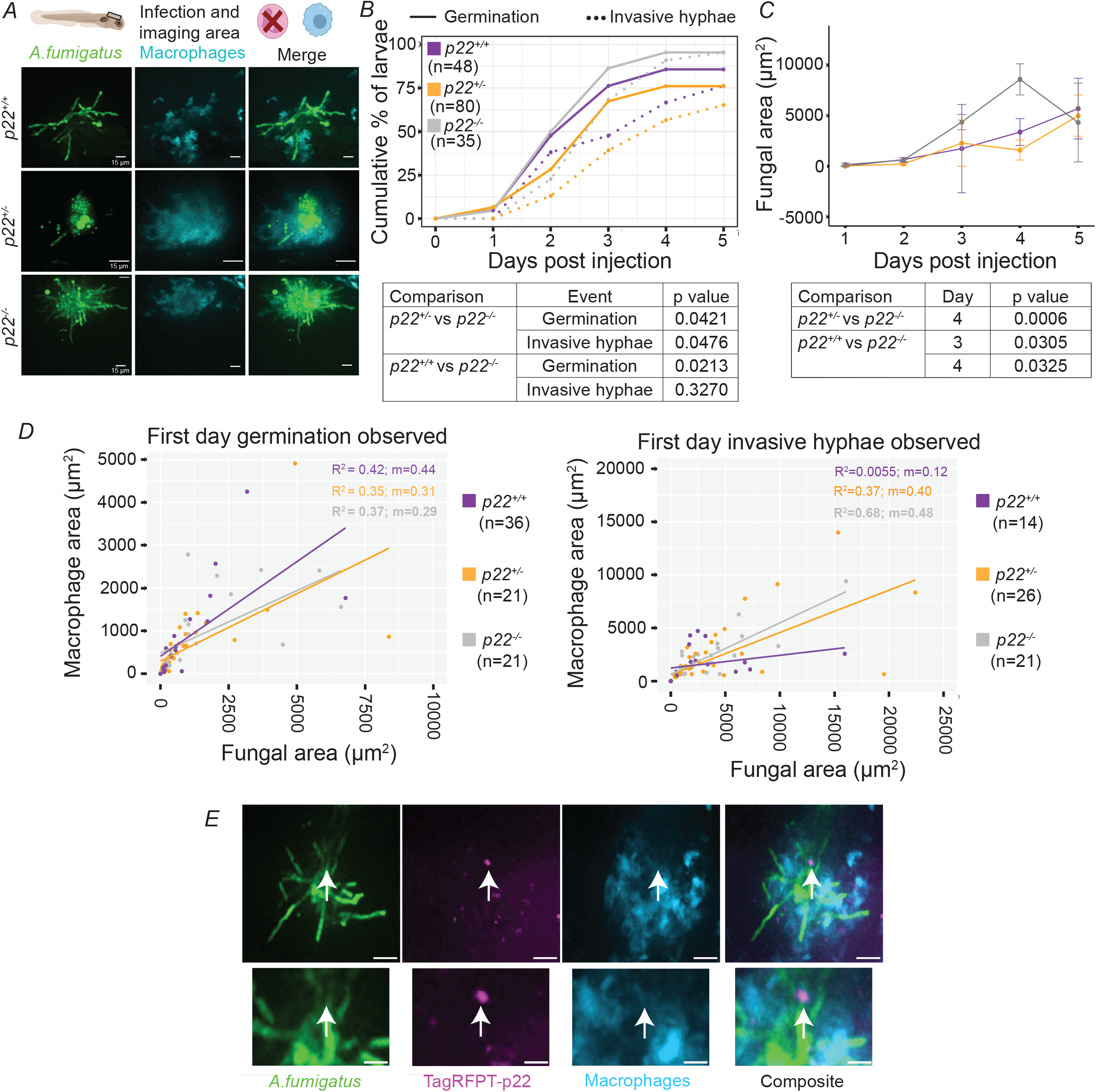
PHOX activity promotes macrophage control of *A. fumigatus* germination and invasive hyphal growth. **A-D.** Neutrophil-defective (*mpx:rac2D57N*) larvae with a stable mutation in *p22* and heterozygous and wild-type sibling controls were infected with the YFP-expressing *A. fumigatus* strain TBK1.1, and imaged daily from 1-5 dpi. Data are pooled from six independent replicates, with a total N of 163. (A) Representative images showing fungal growth and macrophage recruitment at 3 dpi. Images for *p22^+/+^* and *p22^−/−^* larvae were captured at 20× magnification, whereas the representative image for the *p22^+/−^* larva was acquired at 40× magnification. Scale bars, 15 µm. (B) Cumulative percentage of larvae exhibiting fungal germination or invasive hyphal growth over 5 dpi. (C) Quantification of fungal area. Estimated marginal means (emmeans) ± SEM are shown. (D) Correlation between fungal area and macrophage recruitment to the infection site in individual larvae. Measurements were taken on the day when germination (left) or invasive hyphal growth (right) was first observed. Each dot represents an individual larva from six independent replicates. Linear regression with slope and R² values are indicated for each genotype. **E.** Representative image from experiment in Figure 2A-D of macrophage TagRFPT-p22 co-localization with *A. fumigatus* hyphae. Arrow indicates co-localization. Scale bar, 20 µm.

Germination and invasive hyphal growth (Figure 3) were assessed from maximum-intensity Z-projections of Z-stacks containing the infection site. Germination was defined by the emergence of a germ tube from a swollen conidium, and invasive hyphal growth was defined by the presence of branching, elongated hyphae extending beyond the initial site of germination (Thrikawala et al., 2022).These events were manually quantified within the defined ROI of the infection site.

### Statistical Analysis

All statistical analyses were performed in RStudio version 2022.07.2+576 (Posit Software, PBC). Data from independent replicates were pooled for analysis, and total *N* values are indicated in the corresponding figure panels and/or legends.

For larval survival, data from three replicates, with at least 8 larvae per condition per replicate, were analyzed using a Cox proportional hazards model with replicate included in the model to account for variation between independent experiments. Hazard ratios and corresponding *P* values were calculated from the fitted models.

For ROS staining, data from three replicates, with at least 8 larvae per condition per replicate, were analyzed using a linear model. Estimated marginal means (EMMs) were calculated for each experimental condition, and pairwise comparisons were performed using Tukey-adjusted contrasts. Standard errors (SEs) were calculated from the model for statistical inference, and EMMs ± SEM were calculated from the model and EMMs for graphical presentation.

For CFU analysis, counts were normalized to the initial injection dose for the corresponding experimental condition. A linear model including experimental replicate, condition, time, and the condition × time interaction was fitted. EMMs were calculated for each condition at each time point, and pairwise comparisons between conditions within each time point were performed using Tukey-adjusted contrasts. EMMs ± SEM were calculated from the model displayed in graphs.

For co-localization analysis, data from three replicates, with at least 8 larvae per condition per replicate, were analyzed at each time point using a linear model that included experimental replicate. For Figure 2, larval genotype was included as the predictor, whereas for Figure 4, fungal genotype was included as the predictor. EMMs and SEs were calculated from the models and are displayed in the graphs. Pairwise comparisons were performed using Tukey-adjusted contrasts.

**Figure 4.**
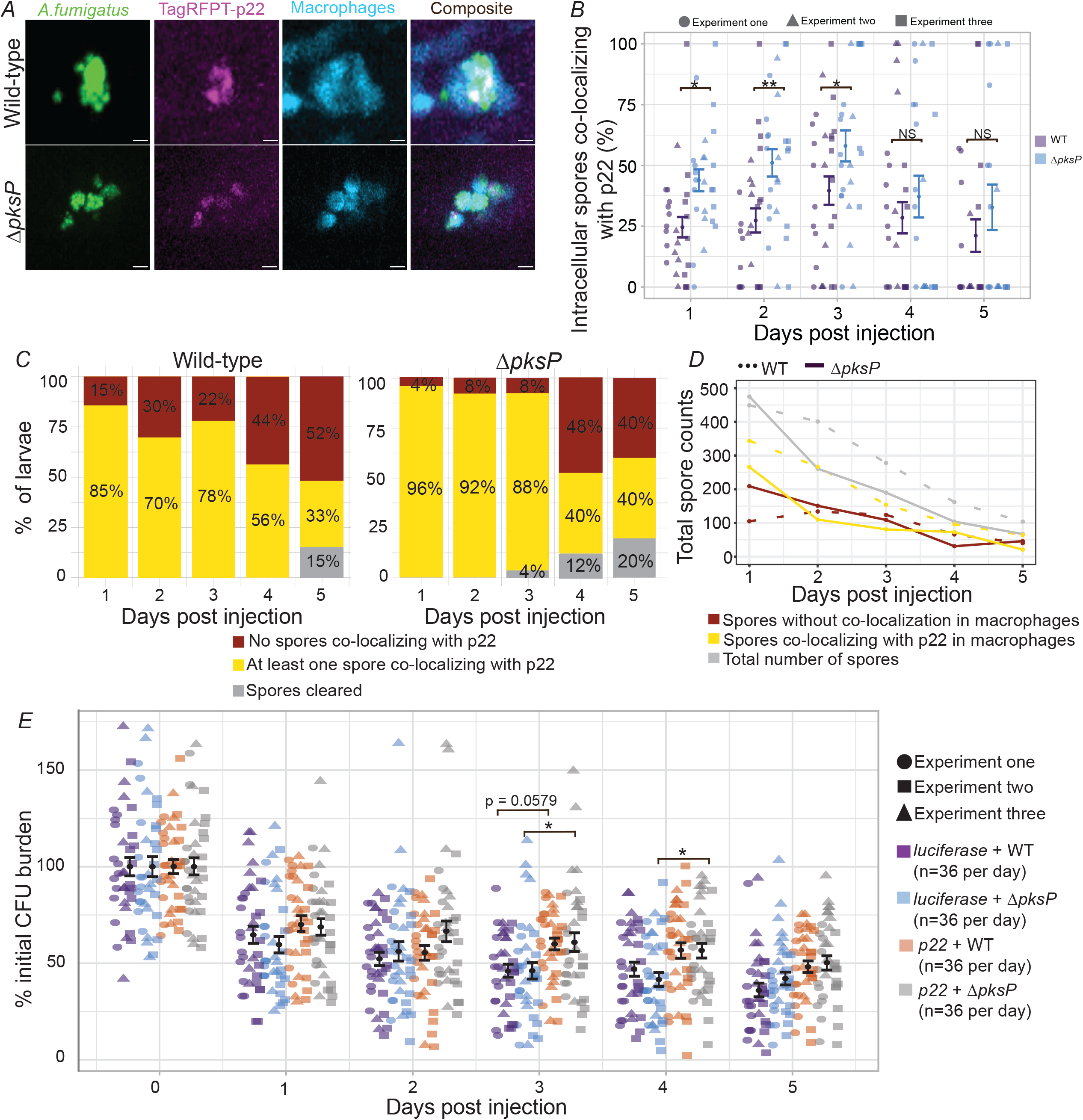
Spores lacking PksP-produced melanin are specifically susceptible to ROS-mediated killing. **A-D.***mfap4:BFP* (macrophage-labeled) zebrafish were crossed with *mfap4:TagRFPT-p22* (p22-labeled) zebrafish. Resulting larvae were infected with either an average of 19 spores of the YFP-expressing *A. fumigatus* strain TBK1.1 or an average of 19 spores of a Δ*pksP* strain derived from TBK1.1. Larvae were imaged daily from 1-5 dpi. Data are pooled from three independent replicates with a total N of 52 larvae (A) Representative images of TagRFPT-p22 (magenta) co-localization with *A. fumigatus* spores (green) within macrophages (blue) at 1 dpi. Scale bar, 10 µm. (B) Quantification of the percentage of spores inside of macrophages that co-localize with p22, with an average of ∼19 spores analyzed per larva. Each dot represents an individual larva. Bars represent estimated marginal means (emmeans) ± SEM. (C) Quantification of the percentage of larvae containing at least one spore co-localizing with p22 within macrophages at each dpi. (D) Total spore counts in different categories across all larvae. **E.** *p22* crispants or *luc* control larvae were generated, infected with TBK1.1 or TBK1.1 ΔpksP strains of *A. fumigatus*, and homogenized for CFU counts from 0–5 dpi, with a total *N* of 36 larvae per condition per day. CFU counts were normalized to the average injection dose at 0 dpi. Each point represents an individual larva, with point shape indicating replicate. Data represent three independent experiments, with 36 total larvae per dpi per condition, with bars indicating estimated marginal means (emmeans) ± SEM.

For imaging analyses of fungal burden, data from six replicates, with at least 8 larvae per condition per replicate, were pooled for analysis. Fungal area was analyzed using a linear model that included experimental replicate, and pairwise comparisons between genotypes were performed using Tukey-adjusted contrasts. EMMs +/-SEM were calculated from the model for graphical presentation. Germination and invasive hyphal growth were analyzed using Cox proportional hazards models that included experimental replicate. Hazard ratios and corresponding P values were calculated from the fitted models.

For imaging analyses of macrophage area, data from six replicates, with at least 8 larvae per condition per replicate, were pooled for analysis. The relationship between fungal burden and macrophage area at the infection site was assessed using a linear regression model that included experimental replicate. The coefficient of determination (R²) and regression equation were calculated from the fitted model.

Statistical significance was defined as P < 0.05. Significance is indicated as P < 0.05 (*), P < 0.01 (**), P < 0.001 (***), and P < 0.0001 (****).

## Results

### ROS promotes macrophage-mediated protection against *A. fumigatus* infection in larval zebrafish

To investigate how the PHOX complex contributes to host defense specifically in macrophages during *Aspergillus fumigatus* infection *in vivo*, we genetically and chemically targeted this complex in a transgenic neutrophil-defective larval zebrafish line (*mpx:rac2D57N*) which relies solely on macrophage function for immunity (Deng et al., 2011). In neutrophil-defective larvae also carrying a stable mutation in the gene encoding the p22 component of the PHOX complex (also known as *cyba*), *p22*^-/-^larvae exhibited significantly reduced survival after infection compared with wild-type or heterozygous siblings, with increased mortality primarily occurring from 3–4 days post injection (dpi) onward (Figure 1A). To confirm these results, we also used CRISPR/Cas9 to generate F0 *p22* crispants in neutrophil-defective larvae (Fig S1), which allowed us to perform experiments with higher Ns. *p22* crispants also had significantly reduced survival after infection compared with *luciferase* gRNA-injected control larvae (Figure 1B). Treatment of *luciferase* control neutrophil-defective larvae with N-acetyl cysteine (NAC), a ROS scavenger, also significantly reduced survival compared to treatment with a PBS control, phenocopying the survival of *p22* crispants (Figure 1C), further supporting the role of ROS in macrophage-mediated host protection. NAC treatment did not further decrease survival of *p22* crispants, confirming that these larvae are PHOX-deficient (Figure 1C). Collectively, these findings suggest that in larvae that rely solely on macrophages to control *A. fumigatus* infection, PHOX-mediated ROS production promotes host survival.

To confirm that *A. fumigatus* infection induces ROS production by macrophages, we injected the ROS-sensitive dye H_2_DCFDA at 2 dpi in neutrophil-defective larvae and quantified dye fluorescence. Infection induced a robust increase in ROS signal in control larvae (Figure 1D–E). This increase was abrogated by NAC treatment and in *p22* crispants, demonstrating the specificity of this staining and the efficacy of both of these conditions in diminishing ROS production (Figure 1D–E). Together, these data demonstrate that macrophages produce ROS through the PHOX complex to promote control of *A. fumigatus* infection.

### *p22* co-localizes with spores in macrophages at a low level

We next began investigating which macrophage functions require the PHOX complex and ROS production. Initially, macrophages phagocytose *A. fumigatus* spores, and the PHOX complex can be recruited to phagosomes to target these spores (Naish et al., 2023; Philippe et al., 2003). We therefore first investigated the recruitment of p22 to fungal spores inside of macrophages in larval zebrafish. We generated a new transgenic zebrafish line which expresses p22 fluorescently-tagged with TagRFPT under a macrophage-specific promoter (*mfap4:TagRFPT-p22*). We crossed this line with a line expressing cytoplasmic BFP in macrophages (*mfap4:BFP*), injected YFP-expressing *A. fumigatus* spores into the resulting larvae, and tracked p22 localization relative to internalized spores by daily confocal imaging. Through this imaging, we could observe instances of co-localization of p22 with internalized spores within macrophages (Figure 2A). However, quantification across three independent experiments demonstrated that the majority of spores within macrophages did not co-localize with p22 throughout the 5-day infection. On average, approximately 25% of intracellular spores co-localized with p22 from 1 to 3 dpi (Fig. 2B). However, there was a very large range across larvae, with some larvae having >50% of spores inside macrophages with co-localization, while in other larvae, zero spores co-localized with p22 (Fig. 2B).

Therefore, we also analyzed the data in terms of whether any spores within a larva had co-localization with p22. We found that 43% of larvae on 1 dpi had at least one spore co-localizing with p22. This percentage increased slightly over time, with 60% of larvae having at least one co-localized spore at 2 and 4 dpi (Figure 2C). However, this occurred as total spores were also cleared over time, with the proportion of larvae that cleared all spores increasing from 10% at 3 dpi to 17% at 4 and 5 dpi (Figure 2C).

Although the percentage of intracellular spores co-localizing with p22 increased over time, the absolute number of co-localized spores remained relatively constant, while the proportion of spores outside macrophages decreased over time (Figure 2D). The apparent increase in co-localization therefore reflects progressive clearance of total spores rather than increased recruitment of p22. Overall, these results demonstrate that p22 can co-localize with spores inside of macrophages in zebrafish, but that the percentage of intracellular spores co-localized with p22 at a given time is relatively low.

### PHOX activity is not required for spore killing

Since we find that p22 can co-localize with spores, but only at low rates, we next wondered whether p22, and therefore phagocyte ROS production, is required for spore killing. To specifically measure spore killing, we infected larvae with a *ΔpyrG* strain of *A. fumigatus,* which requires uridine and uracil supplementation for growth, and cannot efficiently germinate inside of larval zebrafish (Rosowski et al., 2018), and then homogenized larvae for CFU counts over a 5 day infection. In *luciferase* control larvae, spore numbers decreased significantly over the course of infection, demonstrating slow but effective clearance by the host, mirroring previous observations in the larval zebrafish model (Knox et al., 2014; Rosowski et al., 2018) (Figure 2E). Comparison of spore clearance between *luciferase* control and *p22* crispant larvae revealed no statistically significant difference in spore burden at any time point, but with a trend toward increased burden in *p22* crispant larvae at 1 dpi (p = 0.0536) (Figure 2E). These findings indicate that p22 is not required for overall clearance of fungal spores and suggest that the protective effect of p22 on macrophage-mediated host survival is primarily through mechanisms other than spore killing.

### PHOX activity promotes macrophage control of *A. fumigatus* germination and invasive hyphal growth

As PHOX-dependent ROS production is not required for macrophage-mediated spore clearance, we next asked whether the PHOX complex instead inhibits fungal developmental progression during infection. Macrophages restrict infection not only by killing spores but also by limiting germination and subsequent hyphal growth (Rosowski et al., 2018; Tanner & Rosowski, 2024). To test the role of p22 and the PHOX complex in these processes, we infected zebrafish larvae from a p22 stable mutant line with fluorescently-labeled macrophages (*mfap4:BFP*) crossed into the neutrophil-defective background (*mpx:rac2D57N*) with YFP-expressing *A. fumigatus* spores and performed *in vivo* imaging to monitor and quantify fungal germination and invasive hyphal growth over time within the host (Figure 3). In all p22 genotypes, we observed fungal germination and hyphal growth, however, by 3 dpi, *p22^-/-^*larvae exhibited markedly increased germination and extensive hyphal development compared with *p22^+/+^* and *p22^+/-^* controls (Figure 3A). Analysis over 5 dpi revealed that loss of p22 significantly increased the proportion of larvae undergoing fungal germination and invasive hyphal growth (Figure 3B). Specifically, *p22^-/-^* larvae experienced higher rates of germination and hyphal development compared with *p22^+/-^* larvae (p = 0.0421 and p = 0.0476, respectively). Compared with wild-type larvae, *p22* mutants showed a significant increase in germination (p = 0.0213), with a similar but not statistically significant trend toward increased hyphal growth (p = 0.3270). Consistent with increased germination and filamentation, quantification of fungal area demonstrated that *p22^-/-^* larvae exhibit significantly greater fungal growth compared with both *p22^+/+^*and *p22^+/-^* larvae at multiple time points, including at 4 dpi (p = 0.0325 and p = 0.0006, respectively) (Figure 3C). These results indicate that p22 plays a key role in macrophage-mediated restriction of the transition from conidia to invasive fungal morphologies and fungal growth.

To determine whether the increased fungal growth observed in p22-deficient larvae was due to impaired macrophage recruitment rather than a functional defect, we quantified macrophage accumulation relative to fungal burden at the onset of germination and hyphal growth. Across all genotypes, macrophage recruitment, measured as macrophage area at the infection site, scaled proportionally with fungal area (Figure 3D). These results demonstrate that loss of p22 does not affect macrophage migration or accumulation at infection sites, but instead reflects a defect in macrophage antifungal effector function.

As these data suggest that p22 and PHOX activity plays a role in macrophage-mediated control of extracellular hyphal growth, we wondered whether we could observe co-localization of our fluorescently-tagged p22 line with hyphae. Re-examining images from that daily imaging experiment (Figure 2A-D), we identified examples of macrophage-expressed p22 association with hyphal structures (Figure 3E). These images further support the idea that macrophage p22-dependent mechanisms can target later stages of invasive growth.

### PksP-produced melanin inhibits p22 co-localization with spores in macrophages

Fungal melanin is a key virulence factor in *A. fumigatus* that can protect conidia from oxidative damage and interfere with host ROS-dependent antifungal responses (Akoumianaki et al., 2016; Jahn et al., 2000; Sugareva et al., 2006). However, subsequent studies have questioned whether PksP-produced melanin directly protects conidia from exogenous ROS, suggesting that the effects of melanin on oxidative stress may depend on the specific experimental or host context (Keizer et al., 2022). Previous studies have also suggested that PksP-produced melanin can interfere with multiple macrophage antimicrobial pathways, including inhibiting PHOX-dependent activation of LC3-associated phagocytosis by excluding the p22 subunit from the phagosome and by sequestering phagosomal calcium to disrupt calcium–calmodulin signaling (Akoumianaki et al., 2016; Kyrmizi et al., 2013). These findings are consistent with the limited co-localization of p22 with melanized conidia that we observed (Fig. 2A–D).

However, as we also found that loss of macrophage p22 resulted in only limited effects on conidial killing, with larger effects on control of germination and hyphal growth, the relevance of melanin-mediated inhibition of PHOX to fungal infection outcome *in vivo* is still unclear.

We first determined whether melanin affects PHOX recruitment to the phagosome in macrophages in larval zebrafish by comparing p22 co-localization with YFP-expressing melanin-deficient Δ*pksP* and parental wild-type spores over 5 days of infection. Again, we could observe instances of both wild-type and Δ*pksP* spores co-localized with TagRFPT-tagged p22 inside of macrophages (Figure 4A). Focusing only on spores inside of macrophages, quantification of the percentage of spores that co-localized with p22 demonstrated that Δ*pksP* spores showed higher levels of p22 co-localization compared with wild-type spores (Figure 4B). These differences are statistically significant 1-3 dpi, with ∼50% of *ΔpksP* spores co-localized with p22 compared with ∼25% to 30% of wild-type spores. Again, there was substantial heterogeneity in p22-spore co-localization between individual larvae. We therefore also categorized each larva based on whether any infecting spores co-localized with p22 (Figure 4C). This analysis also demonstrated higher co-localization of *ΔpksP* infections with p22, as 88-96% of larvae infected with *ΔpksP* spores had at least one spore co-localizing with p22 in macrophages from 1-3 dpi, compared to 78-85% of larvae infected with wild-type spores. Looking at the total spores across all larvae and all replicates, we again found that, for both infection groups, spores were cleared over time, and therefore the percentage of spores with co-localization is a function of both active co-localization and spore clearance (Figure 4D). Overall, these data suggest that PksP, through melanin production, inhibits engagement of p22 with macrophage phagosomes containing spores, providing *in vivo* confirmation of previous results in cell culture (Akoumianaki et al., 2016; Jahn et al., 2002).

### PksP-produced melanin protects against ROS-mediated spore killing

Since PksP-produced melanin interferes with localization of p22, and therefore likely the PHOX complex, to spores inside of macrophages, we next asked whether PksP affects the resistance of spores to host killing. This idea was also supported by the difference in the total spore counts we measured from the prior imaging analysis, suggesting that Δ*pksP* spores may be cleared more quickly that wild-type spores (Fig 4D). To directly measure spore clearance, we performed CFU analyses on larvae infected with Δ*pksP* or parental wild-type spores over a 5 day infection. To additionally test the interplay between PksP and ROS production, we additionally performed this experiment in both *p22* crispant and *luciferase* gRNA-injected control larvae. Consistent with our previous results (Figure 2E), loss of p22 does not have a major effect on clearance of wild-type spores, with only minor effects at 3 dpi (Figure 4E, purple versus orange, p=0.0579). Surprisingly, clearance of Δ*pksP* spores was also not significantly different than clearance of wild-type spores at any day in *luciferase* control larvae (Figure 4E, purple versus blue). However, within larvae infected with Δ*pksP* spores, loss of p22 had an effect on spore clearance at all days of infection, with these differences reaching statistical significance at 3 and 4 dpi (Figure 4E, blue versus grey, p=0.0385 and p=0.0333). This result demonstrates that during infection with Δ*pksP* spores, p22 significantly promotes spore clearance. Overall, these data are consistent with the conclusion that macrophages have many effector functions for killing spores, not just production of ROS by the PHOX complex. Therefore, inhibition of the PHOX complex by PksP-produced melanin does not play a major role in protecting spores in wild-type hosts. However, when spores do lack PksP-produced melanin, p22-dependent PHOX activity becomes a major contributor to spore killing. Overall, spore clearance reflects a combination of PHOX-dependent and -independent macrophage functions that have varying importance depending on pathogen characteristics, including the presence of melanin.

### PksP-produced melanin antagonizes macrophage-mediated immunity through inhibition of ROS-dependent mechanisms

As our results thus far indicate that modulation of the PHOX complex by PksP-produced melanin only affects spore killing in specific contexts, we next asked how PksP-produced melanin influences fungal virulence and host survival overall and whether any virulence conferred by PksP-produced melanin depends on macrophage ROS production. We first compared survival between wild-type larvae infected with Δ*pksP* spores and parental wild-type spores, finding that in wild-type hosts, which normally control *A. fumigatus* infection (Knox et al., 2014; Rosowski et al., 2018), loss of PksP does not have an effect (Fig 5A). We therefore moved on to experiments in the neutrophil-defective background (*mpx:rac2D57N*), to again focus exclusively on the function of macrophages, and generated *p22* crispants and control *luciferase* gRNA-injected larvae in this transgenic line. In line with our previous results (Fig 1A, B), disruption of p22 in neutrophil-defective hosts significantly decreased survival after infection with wild-type spores (HR = 4.00, p = 2.16 x 10^-8^). This effect was even stronger after infection with Δ*pksP* spores (HR = 5.88, p = 4.26 x 10^-9^), consistent with the idea that when spores lack PksP-produced melanin, p22-dependent PHOX activity is an even larger contributor to fungal control. Comparing the virulence of Δ*pksP* and wild-type parental spores, in *luciferase* control neutrophil-defective larvae (D57N/luc), infection with Δ*pksP* spores resulted in improved survival compared with wild-type spores, consistent with a role for melanin in fungal virulence, although this difference did not reach statistical significance (HR = 0.56, p = 0.066) (Fig 5A). However, in p22-deficient neutrophil-defective larvae (D57N/p22), survival did not differ significantly between wild-type and Δ*pksP* infection, with Δ*pksP*-infected larvae actually experiencing more death (HR = 1.27, p = 0.255), indicating that the pathogenic effect of fungal melanin depends entirely on macrophage PHOX activity. These data support a model in which the main role of melanin in fungal virulence against macrophage-mediated immunity is antagonism of ROS-dependent defenses.

**Figure 5.**
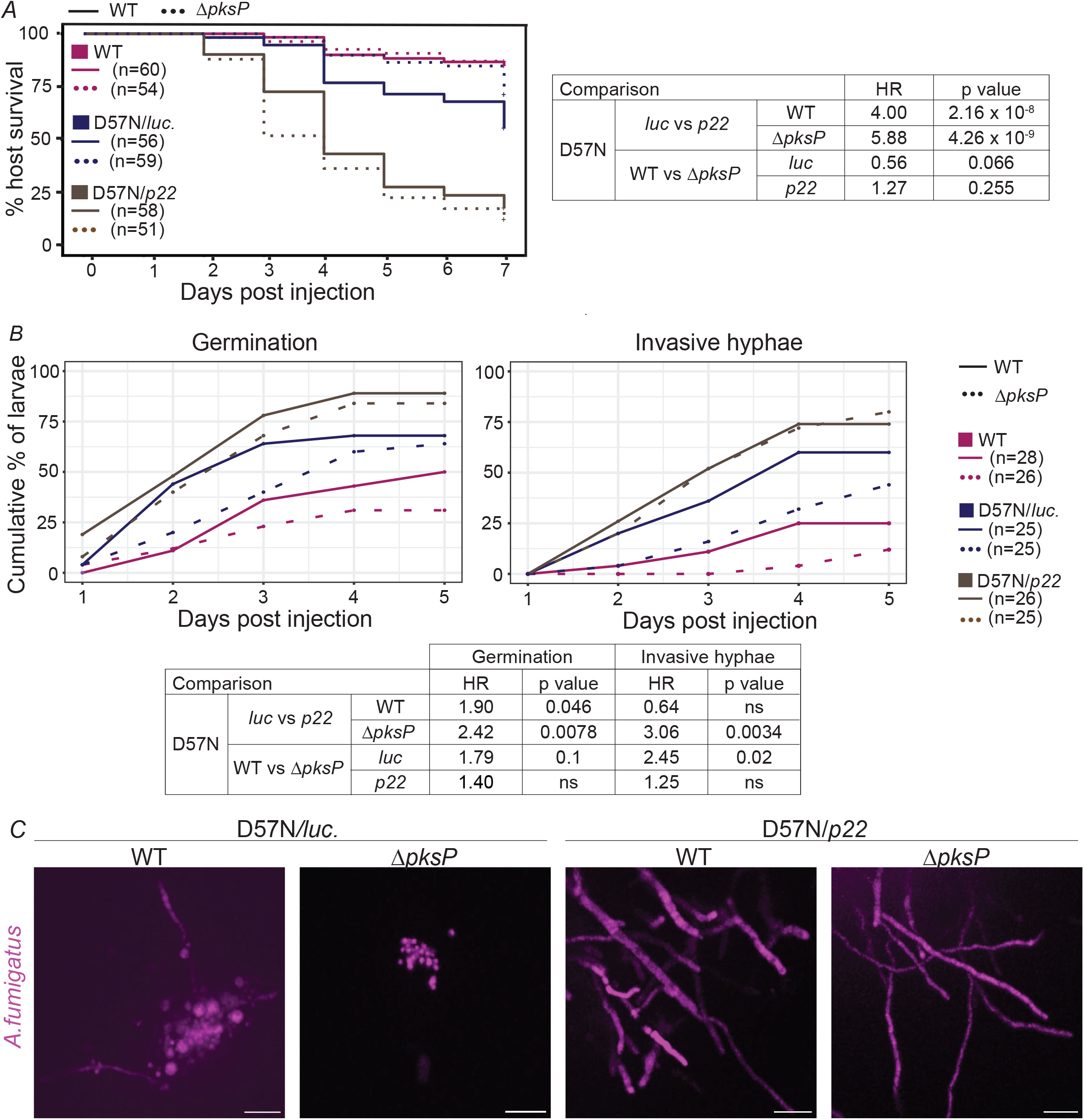
PksP promotes fungal virulence by inhibiting ROS-dependent macrophage control of fungal hyphal growth. **A-C.** Wild-type larvae or *p22 crispants* and *luc* controls in a neutrophil-defective (*mpx:rac2D57N*) background were infected with the RFP-expressing *A. fumigatus* strain TBK 5.1 or the corresponding TBK5.1 Δ*pksP* mutant strain. (A) Survival of larvae was monitored for 7 dpi. The average inoculum was 46 CFUs (WT), 44 CFUs (D57N*/luc*), or 44 CFUs (D57N*/p22)*. Data represent three pooled independent experiments with total N indicated in the figure panel. (B-C) Larvae were imaged daily from 1-5 dpi. Cumulative percentage of larvae exhibiting spore germination or invasive hyphal growth was quantified (B), and representative images of fungal growth are shown at 3dpi (C). Scale bar, 20 µm. Data are pooled from three independent experiments with total N of 155 larvae.

### PksP-produced melanin inhibits macrophage ROS-mediated control of germination and hyphae

We have demonstrated that, (1) the major role for macrophage-generated ROS in controlling *A. fumigatus* infection is through control of germination and hyphal growth (Figure 3), and (2) PksP promotes fungal virulence against macrophages primarily by inhibiting ROS-dependent effects (Fig 5A). These findings suggest that PksP-produced melanin promotes germination and growth of invasive hyphae by inhibiting ROS-dependent macrophage control of these processes. To test this idea, we performed daily imaging of larvae infected with RFP-expressing Δ*pksP* and parental wild-type spores and quantified fungal germination and invasive hyphal formation. Wild-type larvae infected with Δ*pksP* spores experience less germination than when infected with wild-type spores, but this difference was not statistically significant (Figure 5B). We therefore again focused on neutrophil-defective larvae (*mpx:rac2D57N*), generating both *p22* crispants and *luciferase* gRNA-injected controls to test the interaction between PksP-produced melanin and macrophage-produced ROS. Consistent with our earlier experiment (Figure 3), we observed increased appearance of both germination and invasive hyphae in *p22* crispant larvae (D57N/*p22*) compared to *luciferase* control larvae (D57N/*luc*), although this difference was only statistically significant for germination (HR=1.90, p = 0.046) (Figure 5B). Loss of p22 led to an even larger significant increase in observed germination and invasive hyphae after infection with Δ*pksP* spores (germination: HR=2.42, p = 0.0078; invasive hyphae: HR=0.0034, p = 0.0034) (Figure 5B), demonstrating that without the protection mediated by melanin, PHOX-mediated ROS production plays an even larger role in controlling fungal development and growth. By comparing germination and invasive hyphal appearance between larvae infected with wild-type spores and Δ*pksP* spores, we found that wild-type infection leads to significantly more invasive hyphal growth than *ΔpksP* infection in *luciferase* control larvae (D57N/luc: HR = 2.45, p = 0.02), but not in *p22* crispant larvae (D57N/p22: HR = 1.25, p = 0.48) (Fig 5B, C), demonstrating that in the absence of macrophage PHOX-mediated ROS production, PksP-produced melanin is not required for fungal development and growth. Overall, results from this experiment confirm that macrophage-produced ROS helps to control fungal germination and hyphal development and that PksP-produced melanin primarily protects the fungus by countering this ROS and promoting fungal growth.

## Discussion

Macrophage-derived ROS generated by the PHOX complex have been implicated in host defense against *Aspergillus fumigatus*, but the specific function of macrophage PHOX activity to antifungal defense remains unresolved, with studies reporting that macrophages can restrict conidial germination independently of detectable PHOX activity and that macrophages may have a limited role in overall control of infection in some mammalian models (Cornish et al., 2008; Grimm et al., 2011; Mircescu et al., 2009). By combining host and pathogen genetics with longitudinal live imaging in larval zebrafish, we demonstrate that macrophage PHOX activity has distinct stage-specific functions during infection. Our data indicate that PHOX-derived ROS contribute relatively little to killing dormant melanized conidia but are important for restricting fungal germination, invasive hyphal growth, and subsequent disease progression. Thus, fungal developmental progression, rather than conidial killing alone, is a major stage at which macrophage ROS-mediated immunity limits *A. fumigatus* infection.

The limited effect of p22 loss on the killing of wild-type conidia demonstrates that, in larval zebrafish, PHOX is not the principal mechanism responsible for clearing dormant melanized spores. This result contrasts with a previous study demonstrating that ROS production is required for murine alveolar macrophages to kill conidia (Philippe et al., 2003). This discrepancy may reflect differences between in vitro and in vivo models or between host species. In our experiments using neutrophil-deficient hosts, macrophages retained the capacity to limit infection despite the relatively infrequent association of p22 with intracellular wild-type conidia, and loss of p22 had little effect on conidial clearance. These findings indicate that additional macrophage antimicrobial mechanisms contribute to conidial killing, potentially including phagolysosomal acidification and lysosomal antimicrobial activity (Ibrahim-Granet et al., 2003; Latgé & Chamilos, 2019). Because wild-type conidia are coated with PksP-dependent melanin, which can interfere with phagosome maturation and PHOX-dependent antimicrobial pathways, we next considered whether the contribution of PHOX to conidial killing depends on fungal surface composition (Akoumianaki et al., 2016; Chamilos et al., 2016; Jahn et al., 2002). Loss of *pksP* increased p22 association with intracellular conidia, consistent with previous studies showing that PksP-dependent melanin limits PHOX recruitment and ROS-dependent activity at fungus-containing phagosomes (Akoumianaki et al., 2016; Chamilos et al., 2016; Ligeon et al., 2021). However, this increased p22 association did not enhance the overall killing of Δ*pksP* conidia in PHOX-competent hosts. Instead, loss of p22 selectively impaired the clearance of Δ*pksP* conidia, indicating that PHOX becomes an important contributor to conidial killing when melanin-mediated protection is absent, whereas other macrophage mechanisms remain sufficient to control melanized conidia. In contrast, the major contribution of PHOX emerged during fungal development.

Loss of p22 increased fungal germination, growth, and invasive hyphal development, despite relatively preserved killing of wild-type conidia. This finding is consistent with a published study demonstrating that alveolar macrophage PHOX activity limits the germination of inhaled conidia in mice (Idol et al., 2022). Therefore, PHOX-derived ROS is an important mechanism by which macrophages restrict the transition from dormant conidia to invasive fungal growth. This finding extends previous studies demonstrating that macrophages can control *A. fumigatus* through mechanisms beyond direct conidial killing, including inhibition of conidial germination and hyphal development (Rosowski et al., 2018; Tanner & Rosowski, 2024). In particular, we previously demonstrated that macrophages control germination and extracellular hyphal growth through Rac2-dependent mechanisms (Tanner & Rosowski, 2024). Because the phagocyte oxidase complex can incorporate Rac2 (Diebold & Bokoch, 2001), PHOX activity may contribute to the Rac2-dependent control of fungal development. However, Rac2 also regulates other macrophage functions, including transcriptional activation and cell motility, which may additionally contribute to this phenotype (Tanner & Rosowski, 2024).

Our data do not establish where PHOX-derived ROS acts to restrict fungal development. The PHOX complex can function at distinct cellular compartments depending on its site of assembly, thus ROS could act within phagosomes to prevent conidial germination or could be generated at the macrophage surface and act extracellularly on germinating conidia or hyphae. The association of p22 with invasive hyphae is consistent with a potential role for PHOX in this extracellular defense, but does not conclusively establish the site or target of ROS activity. Direct measurements of PHOX activity and ROS dynamics will be needed to distinguish between these possibilities.

The genetic interaction between *pksP* and PHOX indicates that the contribution of fungal melanin to virulence depends on macrophage PHOX activity. A previous study in larval zebrafish suggested that the contribution of fungal melanin to virulence also depends on the *A. fumigatus* strain, with *pksP* deleltion leading to decreased virulence in CEA17 and an International Space Station-isolated strain, but not Af293 (Blachowicz et al., 2020). However, Af293 is the strain background we used and it is unclear why our results differ, demonstrating that *pksP* does promote virulence in Af293. Regardless, our results indicate that any virulence phenotype associated with PksP-dependent melanin is dependent on functional PHOX activity. In PHOX-deficient hosts, deletion of *pksP* no longer reduced fungal virulence, invasive growth, or host mortality. Previous studies demonstrated that *A. fumigatus* melanin inhibits LC3-associated phagocytosis by excluding p22 from fungal-containing phagosomes and that PHOX-derived ROS promote this process through oxidative regulation of ATG4B (Chamilos et al., 2016; Ligeon et al., 2021). Consistent with these findings, we observed increased p22 association with Δ*pksP* conidia *in vivo*; however, we have not yet determined how this impacts LC3 association in our model. Our results extend this model by showing that the melanin–PHOX interaction also influences fungal developmental progression: loss of melanin increased the dependence of conidial clearance on PHOX, while the broader impact of PHOX activity on infection was evident in the restriction of germination and invasive hyphal growth. Thus, PksP-dependent melanin promotes virulence not only by protecting conidia from PHOX-dependent killing, but also by limiting macrophage ROS-dependent control of fungal development.

An important limitation of this study is that our experiments were performed in neutrophil-deficient larval zebrafish, allowing us to isolate macrophage-specific functions but removing the cooperative interactions that occur between macrophages and neutrophils during pulmonary *A. fumigatus* infection. In mammalian models, neutrophils can provide essential protection against germinating conidia and hyphae, and can compensate for loss of alveolar macrophages in some settings (Mircescu et al., 2009). Thus, the contribution of macrophage PHOX activity to fungal development and disease may differ in the presence of functional neutrophils, which could either complement macrophage ROS-dependent restriction or mask some of the phenotypes observed here. In fact, a previous study showed that expression of p22phox solely in neutrophils can be sufficient for control of *Aspergillus nidulans* by larval zebrafish (Schoen et al., 2020). This question is particularly relevant to chronic granulomatous disease (CGD), in which PHOX function is impaired in both neutrophils and macrophages and which is associated with severe susceptibility to *Aspergillus* infections (Marciano et al., 2015; Winkelstein et al., 2000). Our findings suggest that impaired macrophage restriction of fungal developmental progression may contribute to this susceptibility, but the relative contributions of macrophage and neutrophil PHOX activity remain to be determined. Future studies in mammalian models will therefore be important to determine how macrophage PHOX activity integrates with neutrophil and other immune defenses during *A. fumigatus* infection.

In summary, our study identifies a previously underappreciated, stage-specific role for macrophage PHOX-derived ROS during *A. fumigatus* infection. Rather than primarily promoting clearance of dormant melanized conidia, PHOX activity is most important for restricting the transition from conidia to germination and invasive hyphal growth. We further demonstrate that PksP-dependent melanin antagonizes this macrophage defense, linking a fungal virulence factor directly to inhibition of ROS-dependent control of fungal development *in vivo*. These findings shift the focus from conidial killing to developmental progression as a major target of macrophage ROS-mediated immunity and provide new insight into how fungal immune evasion contributes to invasive disease.

## Acknowledgements

We thank members of the Rosowski Lab for helpful discussions and for assistance with zebrafish care. We thank Abigail Godfrey for technical assistance in generation of new Δ*pksP* strains. We thank the Clemson University Aquatic Animal Research Laboratory for providing the microinjection setup used for single-cell embryo injections. This research was supported by a grant from the National Institute of General Medical Sciences of the National Institutes of Health under award number R35GM147464 to E.E.R and the National Institute of Allergy and Infectious Diseases of the National Institutes of Health under award number R21AI164363 to E.E.R. The content is solely the responsibility of the authors and does not necessarily represent the official views of the National Institutes of Health.

## Author contributions

M. Lynette Goins: Conceptualization, Formal analysis, Investigation, Writing - Original Draft

Sourabh Dhingra: Resources

Emily E Rosowski: Conceptualization, Writing - Review & Editing, Supervision, Funding acquisition

## Figure Legends

Supplemental Figure 1. PCR-based assessment of CRISPR/Cas9-mediated excision of the p22 target region. Guide RNAs were designed to introduce double-stranded DNA breaks within exons 4 and 5 of *p22*. The forward primer was positioned upstream of the forward gRNA target site, while the reverse primer was positioned downstream of the reverse gRNA target site. PCR products from five samples injected with p22-targeting gRNAs and samples injected with control luciferase gRNAs are shown. The expected sizes of the intact and excised amplicons are approximately 300 bp and 100 bp, respectively. Bands at approximately 100 bp are visible in the five p22 gRNA-injected samples but not in the luciferase gRNA-injected controls.

